# Can’t decide how much to EAT? Effort variability for reward is associated with cognitive restraint

**DOI:** 10.1101/2020.05.29.123745

**Authors:** Mechteld M. van den Hoek Ostende, Monja P. Neuser, Vanessa Teckentrup, Jennifer Svaldi, Nils B. Kroemer

**Affiliations:** Department of Psychiatry and Psychotherapy, University of Tübingen, Calwerstraße 14, 72076 Tübingen, Germany; Department of Clinical Psychology and Psychotherapy, University of Tübingen, Schleichstraße 4, 72076 Tübingen, Germany

**Author notes:** **Corresponding author**, Dr. Nils B. Kroemer, Calwerstr. 14, 72076 Tübingen, Germany.

## Abstract

Food intake is inherently variable and often characterized by episodical restraint or overeating (uncontrolled eating). Such heightened variability in intake has been associated with higher variability in the brain response to food reward, but it is an open issue whether comparable associations with elevated variability in reward seeking exist. Here, we assessed whether restraint and uncontrolled eating as markers of trait-like variability in eating are associated with higher intra-individual variability in reward seeking as captured by a cost-benefit paradigm. To test this hypothesis, 81 healthy, overnight-fasting participants (*M*_BMI_= 23.0 kg/m^2^ ± 2.95) completed an effort allocation task (EAT) twice. In the EAT, participants had to exert physical effort to earn monetary and food rewards and indicated levels of wanting through visual analog scales (VAS). As predicted, we found that greater trial-by-trial effort variability was associated with lower scores on cognitive restraint, *r*_p_(78) = −.28, *p* = .011 (controlled for average effort). In line with previous findings, higher wanting variability was associated with higher BMI, *r*_p_(78) = .25, *p* = .026 (controlled for average effort). Collectively, our results support the idea that higher variability in reward seeking is a potential risk factor for eating beyond homeostatic need. Since associations with variability measures of reward exceeded associations with average reward seeking, our findings may indicate that variability in the representation of the reward value could be a crucial aspect driving fluctuations in food intake.

## 1. Introduction

Human eating behavior is inherently variable. Intuitively, variability in caloric intake can be found between individuals, as they differ in their metabolic need (Blundell, Finlayson, Gibbons, Caudwell, & Hopkins, 2015), or their tendency to overeat (Davis, 2013). Eating behavior is also variable within individuals (Tarasuk & Beaton, 1991). To illustrate this variability on the intra-individual level, one might consider their favorite Holidays. Many of these celebrations are accompanied by a plethora of special treats or get-togethers that revolve around food. Evidence of indulgences are subsequently found in communal weight gain over holiday seasons, as are indications of subsequent efforts to return to the previous body weight (Helander, Wansink, & Chieh, 2016; Schoeller, 2014; Yanovski et al., 2000). This alternation of indulgence and restriction is not limited to the Holidays. It can be found in our day-to-day eating pattern, which typically consists of overeating in small bouts, using the excess calories during the fasting period until the next meal (Ferrario et al., 2016; Morton, Cummings, Baskin, Barsh, & Schwartz, 2006). These distinct phases introduce variability in food intake. How much we eat is guided by energy homeostasis, which is modulated by an interplay of various internal signals (Ferrario et al., 2016). However, eating behavior can also be motivated by external cues, leading to food intake in the absence of hunger (hedonic eating; (M. R. Lowe & Butryn, 2007)). On the contrary, food intake can also be actively restricted to compensate for previous high caloric intake (Chen, Papies, & Barsalou, 2016). Notably, intra-individual variability in food intake provides a quantification of an individual’s deviation a homeostatic demand, and as such may account for different forms of disordered eating (Neuser, Kühnel, Svaldi, & Kroemer, 2020). To better understand what is driving non-homeostatic eating, it is paramount to understand which traits pertaining to food intake are associated with changes in reward seeking.

In the past decades, functional brain imaging studies have identified altered signaling specific to disordered eating in circuits underlying reward processing and cognitive control. For instance, individuals with anorexia nervosa demonstrate higher cognitive control over reward seeking (Ehrlich et al., 2015; King et al., 2016; Kroemer, 2018). In contrast, studies on individuals with binge eating disorder showed decreased activity in prefrontal regions which are typically associated with cognitive control (Balodis et al., 2013; Hege et al., 2015), whereas increased activity is found in reward circuits (Geliebter, Benson, Pantazatos, Hirsch, & Carnell, 2016; Schienle, Schäfer, Hermann, & Vaitl, 2009; Weygandt, Schaefer, Schienle, & Haynes, 2012). Likewise, it has been demonstrated that greater variability in the brain response to food reward (milkshake receipt) in the nucleus accumbens, a major node of the reward circuit, after a meal is associated with higher BMI and greater variability in subsequent ad libitum caloric intake as well as dietary disinhibition scores (Kroemer, Sun, et al., 2016). It has since been hypothesized that higher intra-individual variability in reward sensitivity is associated with higher variability in food intake (Neuser et al., 2020). One key prediction of the variability theory of food reward is that greater variability in brain signals in the reward circuit would lead to greater variability in reward-related behavior including trait-like food intake and reward seeking. However, it remains to be tested whether variability in reward-related processes such as instrumental behavior or wanting ratings that are commonly used as proxies of food reward signaling are also predictive of inter-individual variability in food intake.

Behavioral tasks that reflect ongoing updates on reward processing are necessary to determine intra-individual fluctuations in reward-related behavior. For example, in cost-benefit paradigms, actions are performed when the benefits outweigh the costs, resulting in a positive utility value (Kroemer, Burrasch, & Hellrung, 2016). Illustratively, consider being offered ice cream as a reward for performing sit-ups. If it is your favorite ice cream, you are likely willing to do more sit-ups compared to a reward of lower value. We are therefore able to use the number of sit-ups as a proxy measure for the ice cream’s reward value. Additionally, you might be more or less tempted by the ice cream, depending on a range of factors (e.g. the weather), leading to differences in how many sit-ups the ice cream is worth to you at a given moment. Thus, fluctuations in the number of sit-ups over time may serve as a measure of variability in the ice cream’s reward value. Analogously, in an effort-based version of cost-benefit paradigms, individuals expend costs in the form of physical effort in order to obtain profits. During the task, the consideration to exert effort at a given point in time is influenced by the signals providing an estimation of the expected benefit and the perceived costs to obtain the reward (Meyniel, Safra, & Pessiglione, 2014; Meyniel, Sergent, Rigoux, Daunizeau, & Pessiglione, 2013; Neuser et al., 2019). According to Meyniel et al. (2013), the costs of effort exertion accumulate over time until the signal surpasses a threshold and effort is suspended. Indeed, work and rest phases are thought to reflect implicit, ongoing updates to the experienced costs (Meyniel et al., 2014; Meyniel et al., 2013). Likewise, updates on the perceived reward value may inform the anticipated benefit of the effort (Kroemer et al., 2014). This is reflected in an increase in effort when higher rewards are at stake (Neuser et al., 2019). Therefore, behavioral variability in instrumental reward seeking may provide an effective means to study the latent variability of value signals in the brain which may help better understand the waxing and waning in food intake.

Markers of variability in eating behavior can be explored through commonly used food questionnaires. For instance, Kroemer, Sun, et al. (2016) were able to show an association between disinhibition and neural variability in response to food reward in a sated state. Scales measuring disinhibition are examples of questionnaires that assess a common core component of eating beyond homeostatic needs, called uncontrolled eating (Vainik, Neseliler, Konstabel, Fellows, & Dagher, 2015). Besides disinhibition, this construct has a variety of indicators, among which are emotional eating and appetitive motivation (Vainik et al., 2015; Vainik, García-García, & Dagher, 2019). By integrating these aspects, uncontrolled eating measures a single dimension that assesses the tendency to overeat (Vainik, García-García, & Dagher, 2019; Vainik et al., 2015). In line with this conceptualization, overeating has previously been described as a scale that reaches from occasional overeating to binge eating symptoms and food addiction (Davis, 2013). Such a scale implies an increasing potential for variability, as more extreme overeating increases the potential range of energy intake as well as the frequency of elevated intake. Furthermore, cognitive restraint has a complementary role in the regulation of eating behavior as it may suppress variability via active control of noise (Manohar et al., 2015). With regard to food intake, it reflects the tendency to actively control eating behavior (Stunkard & Messick, 1985) and could form an underlying construct to uncontrolled eating (Vainik et al., 2019). Thus, cognitive restraint and uncontrolled eating may be complementary indicators of variability in food intake.

All in all, variability in reward processing may provide valuable insight into regular and disordered eating behavior. To test whether aberrant self-reported eating behavior is mirrored in reward-related behavior, we used data from a study investigating the effects of acute transcutaneous auricular vagus nerve stimulation (taVNS) on motivation (Neuser et al., 2019). We used this data because we collected two sessions with the same procedure and anticipated no effects on trial-by-trial variability of the intervention. Variability in reward processes was derived from the intra-individual behavioral variability on an effort task and from intra-individual variability in wanting ratings. We deduced eating behavior from a set of commonly used food questionnaires. In view of previous research (Kroemer, Sun, et al., 2016), we hypothesized that intra-individual variability in instrumental behavior and wanting ratings is associated with measures of variability in eating (i.e., uncontrolled eating and cognitive restraint). Specifically, we expected that behavioral variability can reliably be measured across sessions, as we assume it is a trait-like aspect of behavior. Since stronger incentives have been shown to reduce motor noise (Manohar et al., 2015) we expected lower variability for higher incentives. Additionally, we assessed if differences in variability generalize from food reward to a non-food (monetary) reward. Finally, we expected that behavioral variability is associated with higher uncontrolled eating estimates, whereas we expected lower behavioral variability for higher cognitive restraint.

## 2 Methods

### 2.1 Participants

For the reported analyses, we used the data of 81 participants (47 female, *M*_age_ 25.3 years ± 3.8; *M*_BMI_= 23.0 kg/m^2^ ± 2.95; 17.9 - 30.9) from a study on the effects of acute transcutaneous auricular vagus nerve stimulation (taVNS; (Neuser et al., 2019). Participants were recruited through the University of Tübingen and online channels. We ensured that all participants were right-handed, physically and mentally healthy and had adequate German proficiency to understand the task instructions through a telephone interview. All included participants completed two experimental sessions following the same standardized protocol, once during taVNS and once during sham stimulation. Participants provided written informed consent prior to the first experimental session. They received either money (32€) or course credit in return for their participation in the experiment. In addition to this fixed compensation, participants received money and a breakfast, of which the respective sum and size depended on their task performance. The institutional review board of the Faculty of Medicine, University of Tübingen approved this study (reference number 235/2017BO1), which was conducted in accordance with the ethical code of the World Medical Association (Declaration of Helsinki).

### 2.2 Effort allocation task

To estimate variability in reward processing, we drew on the influence of the expected reward value on effort exertion in an effort-based cost-benefit paradigm. In the effort allocation task (EAT), participants need to allocate their effort in order to optimize wins. The task was adapted from an original paradigm (Meyniel et al., 2013) by changing the input device and adjusting the task conditions. In the adapted version of the EAT (Neuser et al., 2019), participants could earn money and food points by exerting effort in the form of repeatedly pressing a button with their right index finger (button press frequency, FBP).

**Figure 1.**
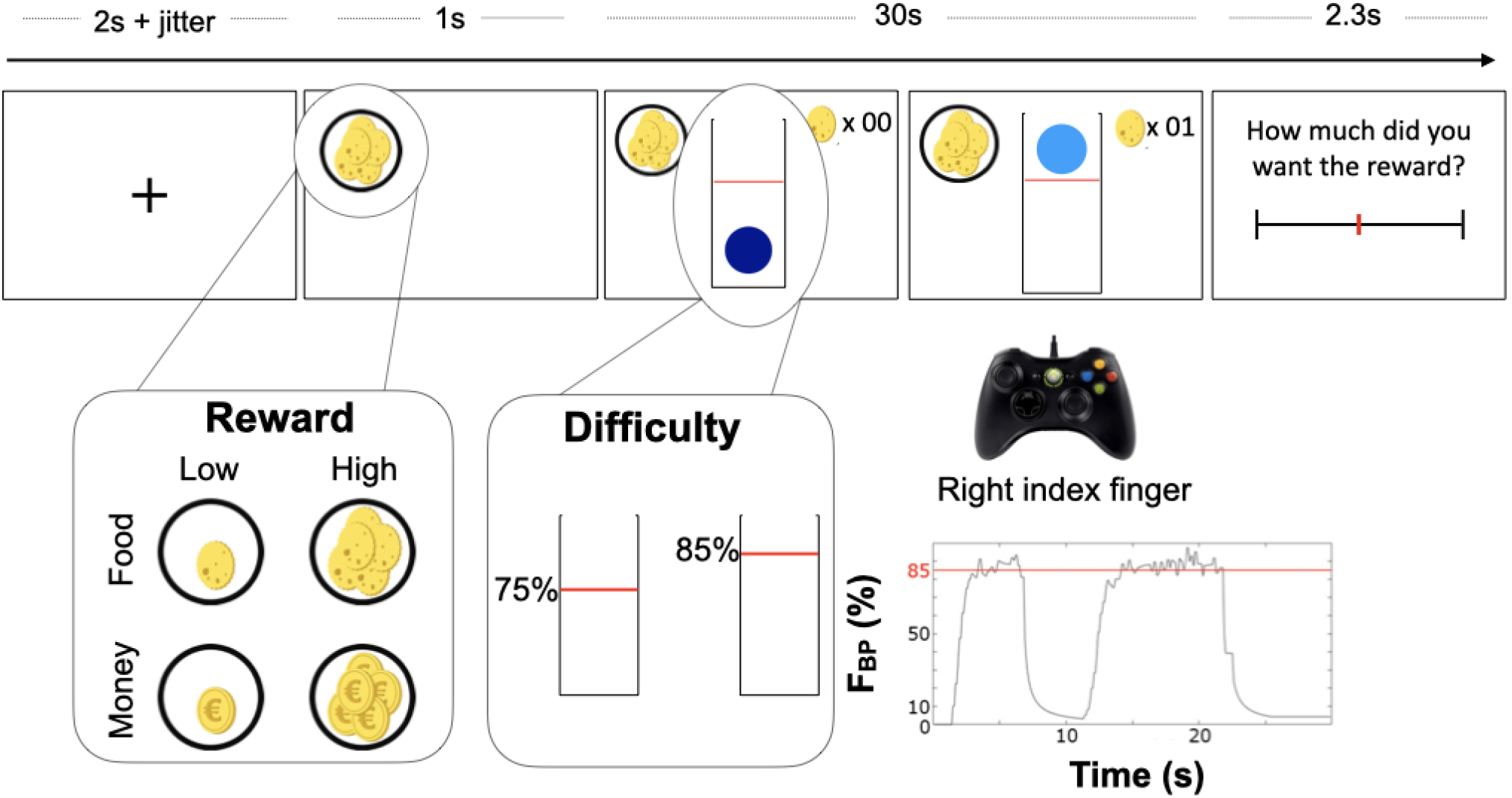
Illustration of the effort allocation task, adapted from Neuser et al. (2019). After a fixation cross, the trial starts with 1 s presentation of the reward cue for that trial. Rewards at stake were depicted by coins for money and cookies for food and also differed in magnitude (low vs. high reward). In the subsequent 30 s effort phase, participants repeatedly press a button with their right index finger to move the ball in vertical direction over a difficulty threshold, indicated by a red line, to collect points. The figure at the bottom right illustrates effort expressed as button press rate (F_BP_) in % of the individual maximum over the course of an exemplary single trial with high difficulty. Finally, participants indicated wanting of the reward at stake on a visual analog scale.

In the EAT, two types of reward were at stake, money (indicated by a coin icon) or food (indicated by a cookie icon). These appeared in one of two magnitudes: low magnitude (indicated by a single icon worth 1 point per second) or high magnitude (indicated by multiple icons worth 10 points per second). In a 30s effort phase, participants could move a ball vertically in a tube by repeatedly pressing a button on an Xbox 360 controller (Microsoft Corporation, Redmond, WA) with their right index finger. Movement of the ball was smoothed to prevent erratic motion. Participants earned points by pressing the button at such frequency that the ball passed above a red line, earning reward points for every full second the ball remained over the line. A reward counter showed the participants their earnings on the current trial. The height of the red line corresponded to the trial’s difficulty, which was either 75% or 85% of the individual maximum FBP as determined during a task training. Difficulty alternated over trials, with the starting difficulty counterbalanced across participants. At the end of every effort phase, participants indicated how much they wanted the reward and how much they exerted themselves on the trial by the means of two visual analog scales (VAS). After the last trial, participants were shown the number of food and money points they had earned during the task. Points were exchanged for a cash payout and calories (in the form of breakfast and a snack) at the end of the session. The exchange rate was set to 1 cent for every 5 money points, and 1 kcal for every 5 food points (Neuser et al., 2019).

The EAT consisted of 48 trials. In the instructions, participants were made aware that the task was too demanding to continuously exert the required effort and therefore, that they could take breaks as needed. In addition, participants had the opportunity to take a short break after half of the trials had been completed. The EAT was displayed using Psychophysics toolbox v3 (Brainard, 1997; Kleiner, Brainard, & Pelli, 2007) in MATLAB v2017a.

### 2.3 Questionnaires

We used the German translation (Strobel, Beauducel, Debener, & Brocke, 2001) of the Behavioral Inhibition System (BIS) and Behavioral Approach System (BAS) scales (Carver & White, 1994) to measure general reward- and punishment-related motivational tendencies. The BIS/BAS questionnaire comprised 24 items that were to be answered on a 4-point Likert scale, ranging from *strong agreement* (1) to *strong disagreement* (4). Although Carver and White (1994) found evidence for 3 subfactors within the BAS items, these factors could not be verified in the German sample (Strobel et al., 2001). Hence, we computed both BIS and BAS scores as an overall sum score of their respective items.

To measure the psychological impact of food abundance in the environment, we used the Power of Food Scale (PFS; (Cappelleri et al., 2009)). The PFS consisted of 15 items, each rated on a 5-point Likert scale, ranging from *I don’t agree (1)* to *I strongly agree* (5). There were three subscales to the PFS, defined by the proximity of the food: available but not directly present in the environment, present but not tasted, and tasted but not consumed. We calculated the score for each subscale by taking an average over its items. All scales of the PFS (partly) reflect uncontrolled eating (Vainik et al., 2019; Vainik et al., 2015).

We used the Three-Factor Eating Questionnaire (TFEQ; (Stunkard & Messick, 1985)) to measure three different aspects of human eating behavior: cognitive restraint, disinhibition and susceptibility to hunger. The TFEQ consisted of 51 items, 21 of which measured cognitive restraint, 16 measured disinhibition and 14 measured hunger. We calculated the factor scores by taking the sum of the items that were applicable to the participant. Out of the three factors, disinhibition and hunger scales are thought to (partly) reflect uncontrolled eating (Vainik et al., 2019). Cognitive restraint might be a separate construct that interacts with uncontrolled eating, as well as body mass index (BMI) and overeating (Vainik et al., 2019).

The Yale Food Addiction Scale (YFAS; (Gearhardt, Corbin, & Brownell, 2009)) measured addiction-like dependence on food. The YFAS consisted of a total of 25 items, sixteen of which were rated on a 5-point Likert scale (ranging from (0) *never* to (4) *4 or more times a week or daily*) and the remaining nine items as *yes (1)* or *no* (0) questions. We calculated the total score by summing over the (recoded) value associated with each answer, in which the Likert scale was valued from 0 to 4, and the binary choice answers were valued either 0 (not applicable) or 1 (applicable) (Meule, Müller, Gearhardt, & Blechert, 2017). The YFAS is thought to be related to more extreme scores of uncontrolled eating (Vainik et al., 2019).

### 2.4 Experimental procedure

Participants completed the questionnaires at home in advance of the first session via SoSciSurvey (Version 2.5.00-i1142; Leiner, 2018) using a personalized link. Participants were invited to two experimental sessions and taVNS vs. sham was applied in a randomized, single-blind crossover design. Each session started in the morning between 7:00 am and 10:15 am and lasted about 2.5 h and participants came to the sessions after fasting overnight (not eating or drinking caloric beverages >8h prior to session). Introductory questions and measurements, among them height and weight, were taken as described by Neuser et al. (2019). Participants subsequently selected their preferred breakfast cereal. We instructed them that they could earn food points and money points during the session. Throughout the sessions, we provided water from which the participant could drink *ad libitum*.

Afterwards, participants responded to the first of three rounds of state questions in the form of a VAS on a computer screen, using the joystick on an Xbox 360 controller. Items focused on metabolic state (hunger, fullness, thirst) as well as mood, as derived from the Positive and Negative Affect Schedule (PANAS; (Watson, Clark, & Tellegen, 1988).

Next, participants completed a practice round of the EAT. Not only did this familiarize the participants with the task, but more importantly, it allowed for an estimation of the individual maximum F_BP_. In two initial trials, an open-ended tube with a blue ball topped with a blue horizontal line was shown on the screen. Participants were instructed to push the line as high as possible by moving the ball. The line stayed at the highest position reached by the ball, even as the button press frequency dropped, indicating the overall maximum FBP. Subsequently, participants completed eight practice trials of the EAT, in which every possible combination of settings occurred once in a randomized order. Crucially, if participants reached a higher maximum FBP on these practice trials, their personal maximum value was updated for the experimental EAT run.

Following the EAT practice, the taVNS electrode was put in place (Neuser et al., 2019). Afterwards, participants completed a food-cue reactivity task (~20 min), the EAT (~40 min), and a reinforcement learning task (Kühnel et al., 2020). Following the last task, participants filled out the state VAS for the second time.

Next, the electrode was removed, and participants received breakfast and a snack in accordance with their food points earned during the EAT. They were given a 10-minute break to eat as much of the cereal as they liked. After the break they reported state VAS for the third time. At the end of each session, participants were given their wins from the completed tasks. Sessions were organized around the same time within a week. An identical standardized protocol was followed for both sessions.

### 2.5 Data analysis

To calculate effort variability, the average relative frequency per trial was determined. Analogous to previous work (Kroemer, Sun, et al., 2016), we subsequently calculated variability as the standard deviation of the model residuals of a linear mixed effects model (Figure 2). For all questionnaires, we used the overall scores of the (sub-) scales for further analyses.

**Figure 2.**
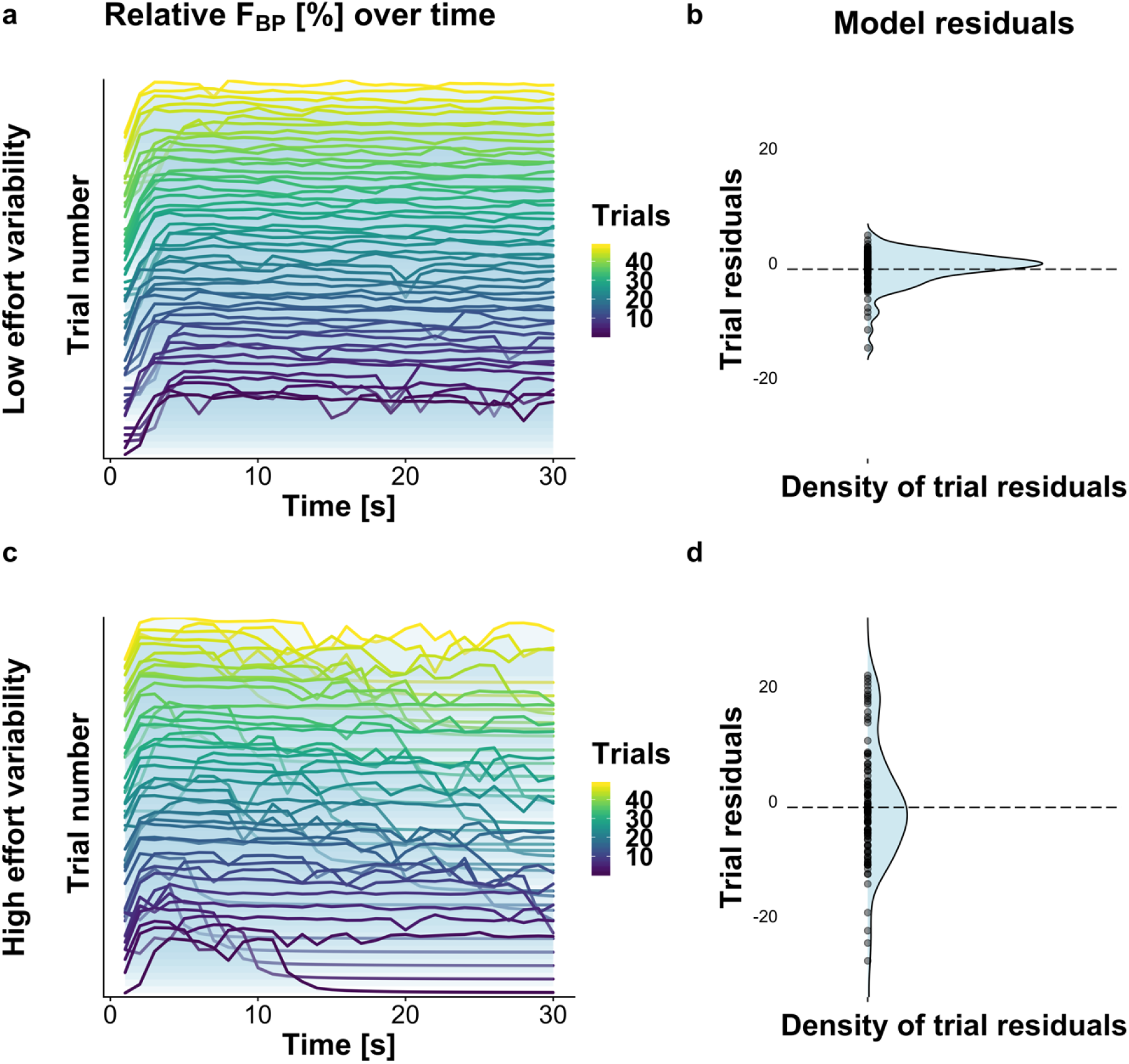
Two illustrative examples of participants, either with high or low effort variability. **A:** Effort in relative button press frequency (F_BP_) in percentage of effort per second (% per s) of the participant with ID 81, demonstrating seemingly consistent behavior. B: Density plot of the trial residuals of participant 81, as calculated through a linear mixed effect model. Each dot under the curve represents the residuals on a single trial. Residuals cluster around zero. Hence, the standard deviation, and therefore the variability estimate of these residuals is relatively low. **C:** Effort in relative F_Bp_ over time of the participant with ID 84. The effort exertion over time appears to be more erratic. D: Density plot of the trial residuals of participant 84. In line with visual estimation, behavior was less predictable to the model, leading to higher residuals. The standard deviation is higher than in B, leading to a higher effort variability estimate for participant 84 than for participant 81.

Next, we tested whether the resulting variability estimates were influenced by taVNS through non-parametric comparison of the stimulation conditions. We then validated that effort variability is trait-like by using the intra-class correlation coefficient (ICC; (Koo & Li, 2016). Providing good test-retest reliability, we averaged variability scores from both sessions to get a better estimation of individual effort variability. With this trait-like variability estimate, we tested for differences in variability between reward magnitudes (low vs. high), as well as differences between reward types (food vs. monetary reward). Analogous to variability, the average effort was estimated by taking the average of both sessions, since stimulation condition did not affect average effort (Neuser et al., 2019).

To test the hypothesized association of heightened effort variability with uncontrolled eating and cognitive restraint, we first examined the Pearson correlations between effort variability, average effort exertion and questionnaire scores. Second, we estimated uncontrolled eating through a confirmatory factor analysis with 2000 bootstrap iterations. For this analysis, uncontrolled eating was included as the latent variable with all questionnaire sub-scales as its indicators, allowing for unique loadings to each of the subscales. Additional correlations between scales were allowed according to modifications indices because our main goal was to optimize the measurement model to estimate associations with uncontrolled eating. Furthermore, effort variability, average effort, and BMI were added as observed variables. We allowed these observed variables to correlate amongst each other and with uncontrolled eating to test our hypotheses.

To verify the associations between effort variability and individual eating-related traits, we also used a tenfold cross-validated elastic net prediction model as a feature selection method. To this end, all questionnaire scores were used as predictors for effort variability, after removal of the shared variance with average effort. As before, this analysis was repeated with average effort instead of effort variability to compare selected features and the accuracy of the prediction.

Finally, to complement the behavioral readout, we analyzed trial-by-trial variability in the ratings of subjective wanting of the rewards at stake in the EAT. Previous research indicated that ratings and effort-based estimates of reward value are closely related with regard to their estimates of the reward value (Lopez-Persem, Rigoux, Bourgeois-Gironde, Daunizeau, & Pessiglione, 2017), as well as bidirectional influences between ratings of a reward and effort to obtain it (Vinckier et al., 2019). Hence, we used wanting variability as a complementary measure of variability in subjective reward value, and wanting variability was added to the existing model as an observed variable to the CFA. In the extended model, correlations between all other observed variables, as well as uncontrolled eating and wanting variability were permitted. Furthermore, a separate elastic net model was used to determine the questionnaires predictive of wanting variability.

Raw data for the analyses were processed and sorted with MATLAB v2019a and R v3.5.2 (R Core Team, 2018). The linear mixed-effects models which provided estimates of effort variability exertion and wanting ratings were implemented in HLM v7 (Raudenbush, Bryk, & Congdon, 2011). The CFAs were performed using AMOS (v26; Arbuckle, 2017) and the elastic net model was implemented in MATLAB using the function lasso (alpha was set a priori to 0.5 providing a balance of lasso and ridge regression). Reliability analysis, as well as correlations and tests of differences of means were performed in IBM SPSS Statistics for Windows, v25.0. A two-tailed *α* ≤ .05 was set as significance threshold for all analyses. Since questionnaire scores and behavioral estimates of reward seeking are correlated within domains, we used cross-validated elastic net models for feature selection which are robust in light of positive correlations among a set of predictors. Figures in the results and data analysis sections were produced with R and AMOS (Arbuckle, 2017).

## 3 Results

### 3.1 Effort variability and average effort are reliable and independent measures

To verify that the variability estimates can be pooled across sessions, we tested whether taVNS had an effect on effort variability. Since we are using residuals, differences in average effort are already accounted for in the statistical model and there was no significant difference in average effort maintenance between taVNS and sham (Neuser et al., 2019). A permutation test with 1000 iterations indicated no difference in effort variability between taVNS (*Mdn* = 6.13) and sham (*Mdn* = 6.39), *p* = .12.

To determine individual consistency of effort variability and average effort, we used the ICC and rank correlation. Both average effort and effort variability demonstrated good to excellent intra-individual consistency between sessions. We found a positive association between average effort in the two sessions, *ρ*(79) = .90, *p* < .001. The *ICC* of average effort was .88, 95% *CI* [.81, .92]. Likewise, we found a positive association between effort variability in the two sessions, *ρ*(79) = .83, *p* < .001. The *ICC* of effort variability was .82, 95% *CI* [.73, .88]. To provide a characteristic estimate of average effort and effort variability, we averaged the estimates for further analyses of inter-individual differences in variability.

**Figure 3.**
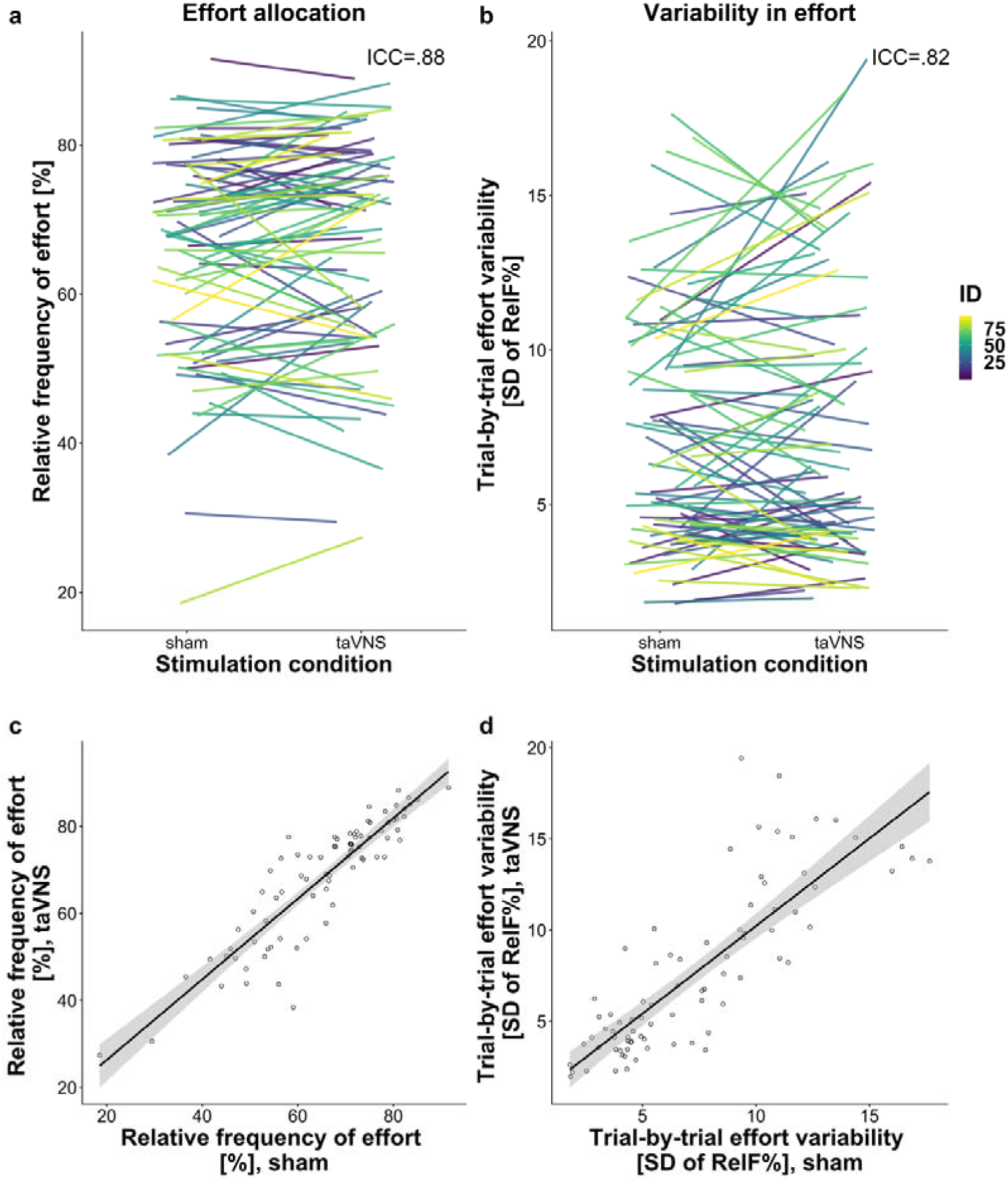
Test-retest reliability of effort variability is comparable to average effort. A: Average effort, relative to the individual’s maximum effort, for both stimulation conditions. Each line represents the change in average effort between sessions of a single participant. Note that order of stimulation conditions was randomized across participants. B: Effort variability, measured as the standard deviation of the residuals when predicting average effort, is plotted analogous to average effort. Colors indicate individual participants. C: Average effort across trials during active transcutaneous auricular vagus nerve stimulation (taVNS) is plotted against average effort across trials during sham. Each dot represents a participant. The trendline represents a simple linear regression between the two conditions, and the shaded region indicates the 95% confidence interval. D: Effort variability across trials during active taVNS is plotted against effort variability across trials during sham stimulation analogous to average effort.

Next, we tested for potential effects of reward type and magnitude on effort variability. A permutation test with 1000 iterations indicated no differences in effort variability between monetary (*Mdn* = 6.24) and food reward (*Mdn* = 6.30), *p* = .35. Since the correlation between the two was high, *ρ*(79) = .93, *p* < .001, it is unlikely that reward type affects the analysis of inter-individual differences in effort variability. Instead, a permutation test with 1000 iterations indicated that effort variability on low reward magnitude trials (*Mdn* = 6.65) was larger than on high reward magnitude trials (*Mdn* = 5.84), *p* < .001. However, due to the high correlation between the two, *ρ*(79) = .72, *p* < .001, this is unlikely to affect the analysis of inter-individual differences in variability. Thus, both average effort and effort variability demonstrated excellent test-retest reliability between two sessions and a plausible modulation by reward magnitude and can therefore be considered as trait-like individual characteristics.

In addition to effort variability, we explored trial-by-trial variability in VAS wanting ratings as a subjective measure of reward evaluation. We therefore assessed whether wanting variability is trait-like and can be used to uncover associations with measures of eating behavior as well. To evaluate whether the stimulation condition had an effect on wanting variability, we first used a permutation test with 1000 iterations. This revealed no significant difference between taVNS (*Mdn* = 8.27) and sham (*Mdn* = 8.57), *p* = .50. Furthermore, wanting variability correlated between sessions, *ρ*(79) = .68, *p* < .001, and its *ICC* was .73, 95% *CI* [.61, .82]. Therefore, test-retest reliability was moderately high, and we obtained a single estimate for each participant’s wanting variability by taking the average value of both sessions. To test whether reward value and reward type affected wanting variability, we used permutation tests with 1000 iterations each. These revealed that wanting variability was higher for small rewards (*Mdn* = 9.72) than large rewards (*Mdn* = 7.65), *p* < .001. Permutation tests also demonstrated that variability in monetary reward (*Mdn* = 8.31) was lower than variability in food rewards (*Mdn* = 8.78), *p* = .015. However, due to the high correlation between the two, *ρ*(79) = .84, *p* < .001, this is unlikely to affect the analysis of inter-individual differences in variability. Thus, we used the average across the reward conditions.

### 3.2 Effort variability correlates with cognitive restraint independent from average effort

To assess associations between effort variability, average effort, individual questionnaire scores, and BMI, we used Pearson correlations. As expected, questionnaires that reflect uncontrolled eating correlated with each other (TFEQ, PFS, and YFAS; Figure 4). Notably, BAS reward sensitivity and TFEQ cognitive restraint demonstrated a rather limited relation to the other questionnaire scores. On the one hand, BIS correlated significantly with both the disinhibition dimension of the TFEQ and the food present subscale from the PFS. On the other hand, BAS had an unexpected significant negative correlation with the YFAS. It further correlated positively with the food tasted subscale of the PFS. Of the questionnaires, only the BIS demonstrated a significant (negative) correlation with BMI although this could be due to the moderate range in our sample.

**Figure 4.**
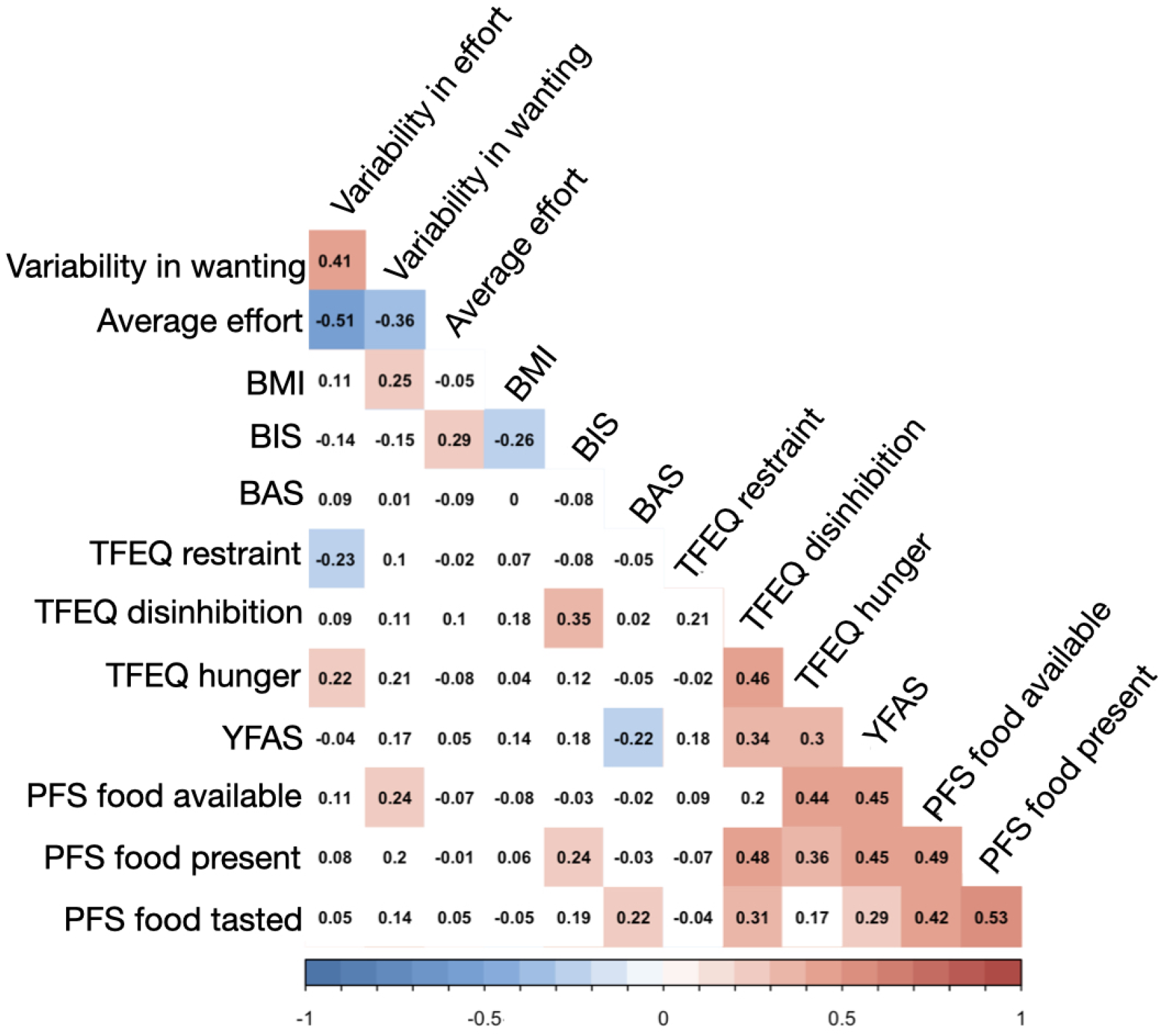
Effort variability correlates with average effort, cognitive restraint and susceptibility of hunger. Correlation coefficient between variables is found in the cell corresponding to the respective row and column. All significant correlations are indicated by color, in which blue represents negative correlations and red positive correlations. Colors follow a gradient, which becomes more saturated as correlation coefficients get higher. BAS = Behavioral Approach System; BIS = Behavioral Inhibition System; BMI = Body Mass Index; PFS = Power of Food Scale; TFEQ = Three Factor Eating Questionnaire; YFAS = Yale Food Addiction Scale.

Effort variability was related to the cognitive restraint and susceptibility to hunger subscales of the TFEQ and average effort (Figure 4). A partial correlation showed that the negative association between effort variability and cognitive restraint remained significant after controlling for the association between variability and average effort, *r*_p_(78) = −.28, *p* = .011. However, the correlation of effort variability with susceptibility to hunger did not remain significant after controlling for average effort, *r*_p_(78) = .21, *p* = .061. In comparison, average effort correlated only with BIS scores (Figure 4), **a** correlation that remained significant after controlling for effort variability, *r*_p_(78) = .26, *p* **=** .021. Wanting variability correlated with effort variability, average effort, PFS food available and BMI. Thus, our results suggest that effort variability shows dissociable associations from average effort (Figure 5) while being associated with wanting variability, a subjective measure of variability in reward evaluation.

**Figure 5:**
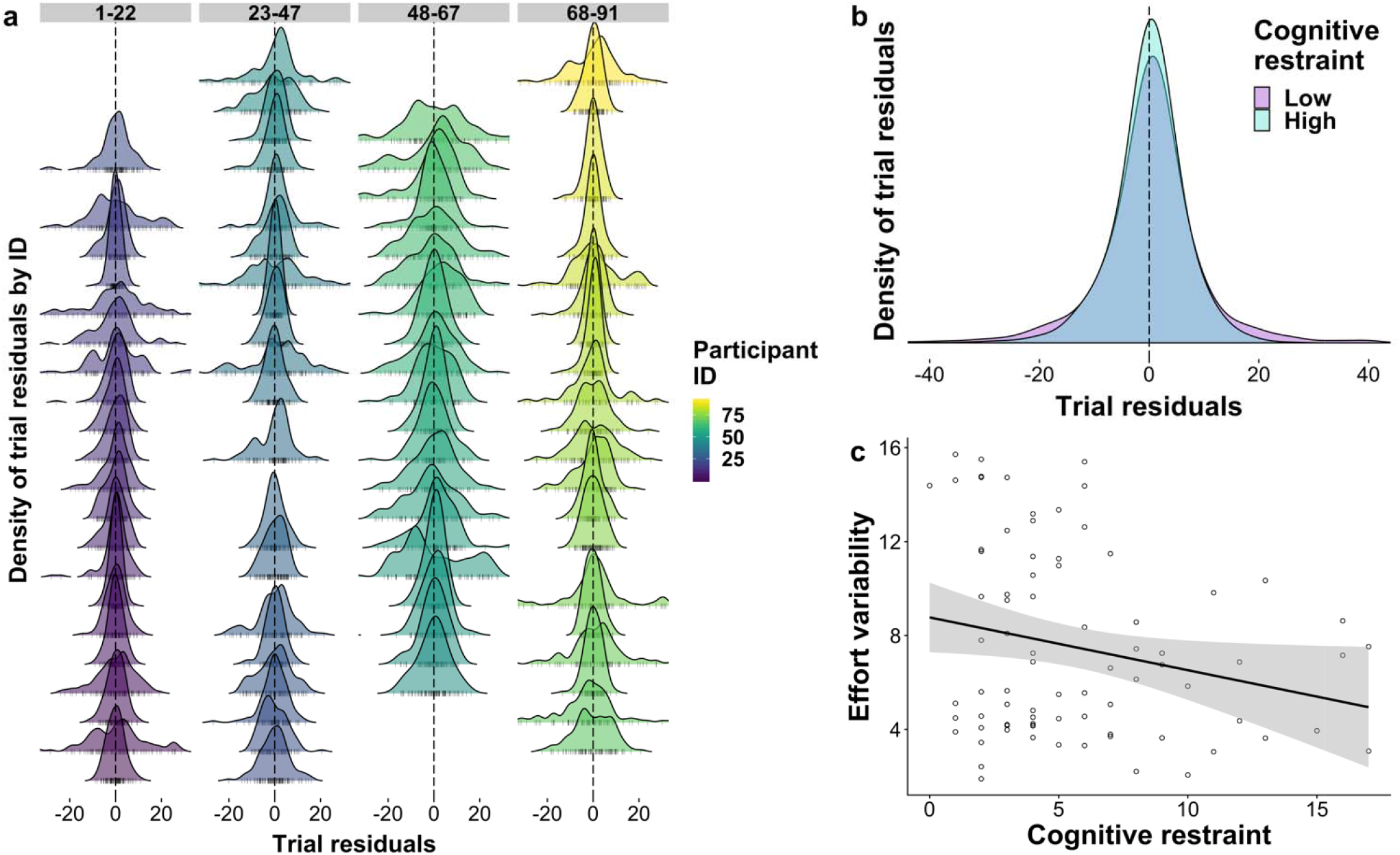
Effort variability differs between individuals and cognitive restraint scores. A: Density of the trial residuals of the linear mixed-effects model for each participant (ID), rows contain binned IDs. B: Trial residuals aggregated for participants with the low (blue, lower quartile: score ≤ 3) and high subscale (yellow, upper quartile: score ≥ 8) scores on the TFEQ cognitive restraint. C: Effort variability plotted against cognitive restraint scores. Each dot represents a participant. The trendline represents the correlation between the two conditions, and the shaded region indicates its 95% confidence interval.

To better evaluate which individual questionnaire scores are associated with effort variability, we used an elastic net model with ten-fold cross validation as feature selection method. As features predicting effort variability after removal of the shared variance with average effort in held-out fold, the model selected cognitive restraint (*b* = - 0.15), susceptibility to hunger (*b* = 0.09) and disinhibition (*b* = 0.08). In contrast, for average effort the elastic net model selected only BIS (*b* = 0.26) as a feature. For wanting variability, the elastic net model selected PFS food available (*b* = 0.57), PFS food present (*b* = 0.17), TFEQ susceptibility to hunger (*b* = 0.08), BIS (*b* = −0.03), and BMI (*b* = 0.17).

**Figure 6:**
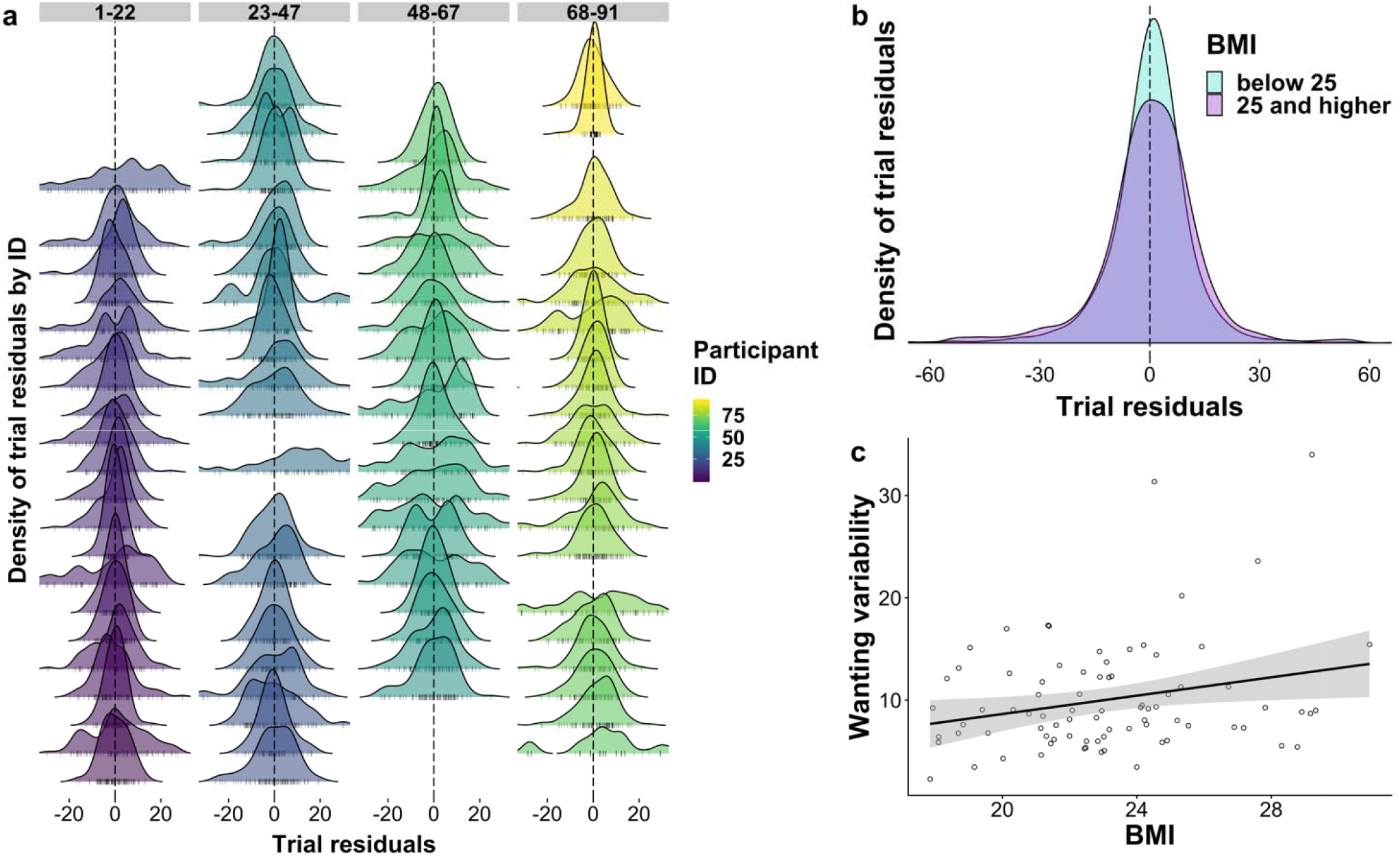
Wanting variability differs between individuals and BMI. A: Density of the trial residuals of the linear mixed-effects model for wanting of each participant (ID), rows contain binned IDs. B: Aggregated trial residuals for low (light blue; BMI < 25 kg/m^2^) and high (dark blue; BMI ≥ 25 kg/m^2^) body mass index (BMI). C: Wanting variability plotted against BMI. Each dot represents a participant. The trendline represents the correlation between the two conditions, and the shaded region indicates its 95% confidence interval.

### 3.3 Effort variability is associated with cognitive control, rather than uncontrolled eating

To estimate uncontrolled eating, we used a CFA in which uncontrolled eating was the latent variable with the individual questionnaires as manifest indicators. Additionally, effort variability, average effort, and BMI were added as observed variables. The initial model without cross-loadings had poor fit (*χ*^2^ = 105.65, *df* = 51, *p* <.001, *CFI* = .704, *RMSEA* = .116), but incorporating correlations among questionnaires in addition to uncontrolled eating improved the model considerably (*χ*^2^ = 70.19, *df* = 45, *p* = .010, *CFI* = .864, *RMSEA* = .084; Figure 7). Moreover, an additional relationship between cognitive restraint and effort variability was indicated by the modification indices and therefore added to the model. These adaptations yielded our final model, which demonstrated reasonable fit (*χ*^2^ = 62.51, *df* = 44, *p* = .034, *CFI* = .900, *RMSEA* = .073). Thus, this model provided no evidence for an association between effort variability in a hungry state and uncontrolled eating, but pointed to an association with cognitive restraint instead.

**Figure 7.**
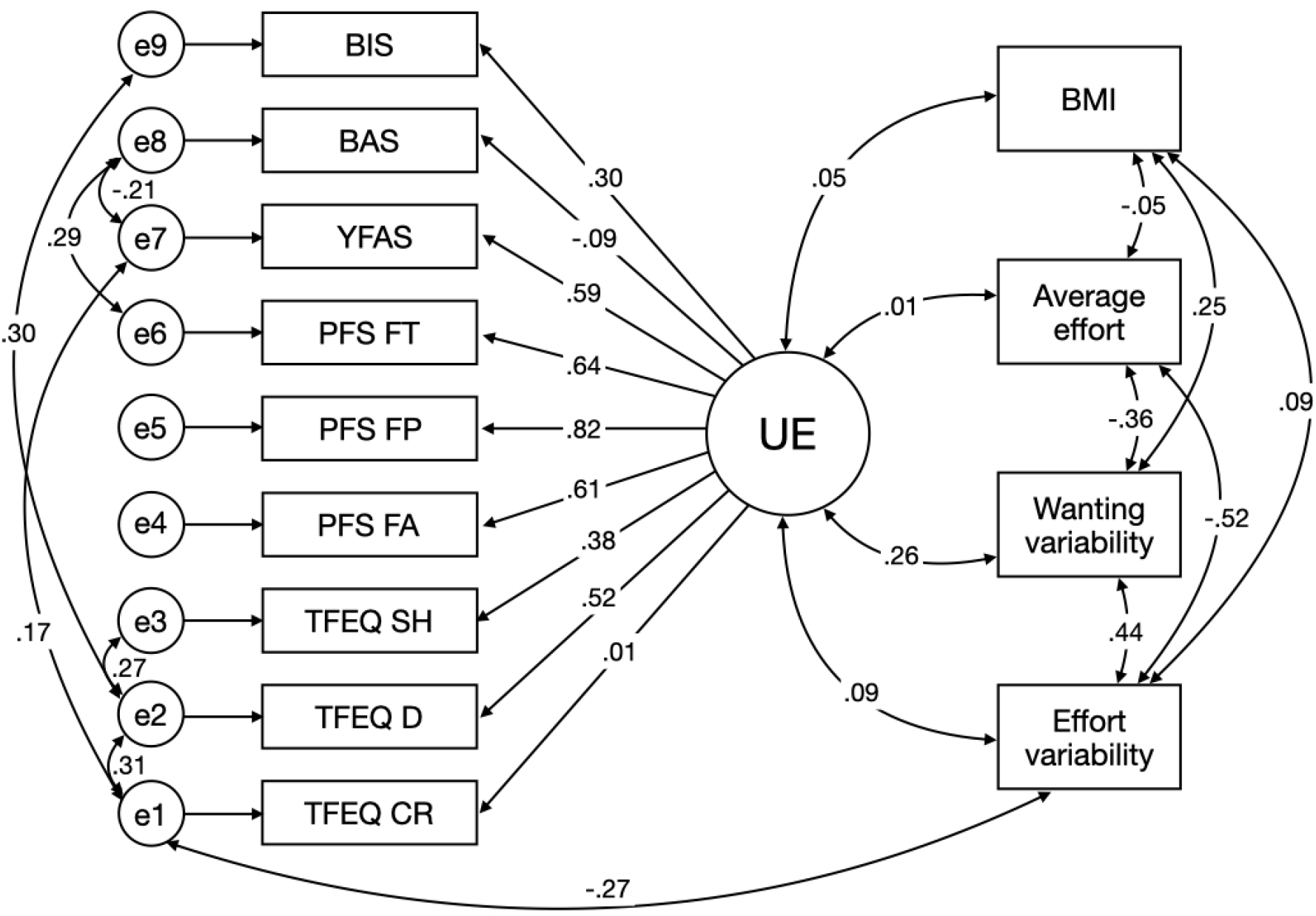
Effort variability is not associated with uncontrolled eating, but with cognitive restraint as shown through confirmatory factor analysis (CFA). Single headed arrows indicate paths, double headed arrows covariances. The initial model did not allow for covariances between the unique loadings e1 through e9. Wanting variability was excluded from earlier model by setting its covariances to zero. BAS = Behavioral Approach System; BIS = Behavioral Inhibition System; BMI = Body Bass Index; PFS FA = Power of Food Scale, Food Available; PFS FP = Power of Food Scale, Food Present, PFS FT = Power of Food Scale, Food Tasted; TFEQ CR = Three Factor Eating Questionnaire, Cognitive Restraint; TFEQ D = Three Factor Eating Questionnaire, Disinhibition; TFEQ SH = Three Factor Eating Questionnaire, Susceptibility to Tunger; UE = Uncontrolled Eating; YFAS = Yale Food Addiction Scale.

To assess the relation between wanting variability and uncontrolled eating, we added it as an observed variable to our structural model (*χ*^2^ = 64.60, *df* = 52, *p* <.113, *CFI* = .938, *RMSEA* = .055; Figure 7), but only found the correlation at a trend level (*r* = .23, *p* = .053). Therefore, larger samples with a greater range in uncontrolled eating might be necessary to provide conclusive evidence on an association. Nevertheless, in the current sample, we were able to demonstrate an association between wanting variability and BMI despite a moderate range of BMI.

## 4 Discussion

Although variability is a central characteristic in eating behavior, its explanatory role in mechanistic theories on the etiology of eating behaviors has long been disregarded as mere noise. Here, using an effort task, we exploited the fact that subjective value of prospective rewards translates to motivation to pursue these rewards. We demonstrated that effort and wanting variability are reliable components of behavior that exhibit relevant ties to habitual food intake. Specifically, we showed that effort variability is negatively correlated with cognitive restraint, whereas wanting variability is positively correlated with BMI. Critically, these associations did not become apparent from an analysis using the average effort, despite correlations between both variability measures, corroborating the theory that variability in reward evaluation is a crucial aspect of eating behavior (Kroemer, Sun, et al., 2016; Neuser et al., 2020). Therefore, our study provides new insights into reward processes driving variability in eating behavior potentially indicating a promising avenue for a better understanding of eating and metabolic disorders.

In our task, we measured subjective reward value via effort and subjective wanting ratings. We found that variability in the reward value measured through instrumental behavior was associated with cognitive restraint independent from other constructs. This negative association with effort variability may be an illustration of the functionality cognitive control has over food intake. Particularly, cognitive restraint, the tendency to exert cognitive control over food intake (Stunkard & Messick, 1985) is a strategy for individuals to confine food intake (Johnson, Pratt, & Wardle, 2012; C. J. Lowe, Reichelt, & Hall, 2019). In accordance with the role of cognitive control as a mediator of food-related impulses (C. J. Lowe et al., 2019), this implies that cognitive restraint may help individuals to limit non-homeostatic overeating. In previous research, Kroemer, Sun, et al. (2016) demonstrated a correlation between disinhibition and variability in food-reward signaling in sated individuals and no significant association was found the same participants in a hungry state. By definition, disinhibition refers to a loss of control over intake and it is conceivable that this should be studied in a sated state instead of a hungry state as in the current study. Future research is necessary to create a more complete understanding of the interaction between metabolic state, trait-like eating behavior and variability in the reward value. Notwithstanding, the current study demonstrates that variability in instrumental behavior on an effort task can provide a low-cost measure of the latent variability in a putative reward value signal therefore adding to the emerging evidence on the relevance of variability in reward.

Crucially, we were able to complement these findings with wanting ratings, which provide a complementary subjective measure of the same latent evaluation process. Although effort exertion and wanting ratings may be closely related (Lopez-Persem et al., 2017; Vinckier et al., 2019), they can be partially dissociated under specific circumstances (Finlayson, King, & Blundell, 2008). In our study, we found unique correlates of both measures, demonstrating some unique component to each of them. For example, individuals with higher wanting variability had also higher BMI values. This can be taken as independent replicate of previous research that demonstrated associations between variability in the brain response to rewards and BMI (Kroemer, Sun, et al., 2016). In light of the current study, wanting variability thus acted as a behavioral proxy for variability in the reward value in hungry participants, similar to effort variability. Identifying such proxies is paramount for future applications, as it provides an accessible gateway to estimate fluctuations in reward value signals guiding behavior.

We identified several limitations of our study that should be considered in future research. First, the current sample consisted of young individuals with a BMI between 17.9 and 30.9 kg/m^2^ who mostly scored low or intermediate on the eating-related questionnaires. Despite the limited range, we were able to identify significant relations between variability and traits pertaining to food intake. However, including participants with a wider range of eating behavior in future studies is a critical next step to solidifying our knowledge of the relationship between variability in reward processes and eating behavior. Second, our estimation of eating behavior was fully based on self-report questionnaires. Future studies could make use of ad libitum meals to provide a lab-based measure of food intake, as has been done previously (Kroemer, Sun, et al., 2016). Third, we identified different markers of variability compared to previous research, which investigated sated rather than hungry participants (Kroemer, Sun, et al., 2016). As such, it is imperative to understand the dynamics of factors influencing variability over different metabolic states. Hence, future research should aim to embed the correlates of variability in the context of metabolic state, be it hungry, sated, or relevant stages in between such as feeling neither hungry nor full. Finally, we used effort and wanting variability as proxies for variability in the reward signal. Complementing this with functional neuroimaging techniques will allow for a direct demonstration of the connection between the measures of variability used here and the underlying neural correlates.

In summary, we demonstrated that intra-individual variability in instrumental behavior, as well as in rated wanting of rewards convey valuable information about inter-individual differences in eating behavior. In contrast to the frequent assumption that trial-by-trial effort or wanting variability is mostly noise, we reported task-based estimates with sufficiently high test-retest reliability for individual prediction (Fröhner, Teckentrup, Smolka, & Kroemer, 2019). Specifically, we showed that individuals with higher cognitive restraint showed lower intra-individual effort variability and that a higher BMI was associated with greater wanting variability in a hungry state. Collectively, these results indicate that typical behavioral measures may help uncover variability in reward processes that is related to eating behavior. Moreover, associations with variability were seen beyond associations with average effort suggesting that they provide incremental information which should be capitalized on in eating-related research. To conclude, we posit that effort and wanting variability may provide a promising avenue to advance our understanding of both healthy and disordered eating.

## Acknowledgements

We thank Caroline Burrasch, Franziska Müller, Sandra Neubert, Moritz Herkner, Magdalena Ferstl, and Leonie Osthof for help with data acquisition. The study was supported by the University of Tübingen, Faculty of Medicine fortune grant #2453-0-0. MMHO, MPN & NBK received support from the Else Kröner-Fresenius Stiftung, grant 2017_A67.

## Author contributions

NBK was responsible for the study concept and design. MPN coded the paradigm and collected data under supervision by NBK. MMHO & NBK conceived the methods and processed the data. MMHO & NBK wrote the draft and all authors contributed to the interpretation of findings, provided critical revision of the manuscript for important intellectual content and approved the final version for publication.

